# Inferring the heritability of large-scale functional networks with a multivariate ACE modeling approach

**DOI:** 10.1101/437335

**Authors:** Fernanda L. Ribeiro, Felipe R. C. dos Santos, João R. Sato, Walter H. L. Pinaya, Claudinei E. Biazoli

## Abstract

Recent evidence suggests that the human functional connectome is stable at different time scales and unique. These characteristics posit the functional connectome not only as an individual marker but also as a powerful discriminatory measure characterized by high intersubject variability. Among distinct sources of intersubject variability, the long-term sources include functional patterns that emerge from genetic factors. Here, we sought to investigate the contribution of additive genetic factors to the variability of functional networks by determining the heritability of the connectivity strength in a multivariate fashion. First, we reproduced and extended the connectome fingerprinting analysis to the identification of twin pairs. Then, we estimated the heritability of functional networks by a multivariate ACE modeling approach with bootstrapping. Twin pairs were identified above chance level using connectome fingerprinting, with monozygotic twin identification accuracy equal to 57.2% on average for whole-brain connectome. Additionally, we found that a visual (0.37), the medial frontal (0.31) and the motor (0.30) functional networks were the most influenced by additive genetic factors. Our findings suggest that genetic factors not only partially determine intersubject variability of the functional connectome, such that twins can be identified using connectome fingerprinting, but also differentially influence connectivity strength in large-scale functional networks.

## Introduction

In the past few years, fMRI research has been living a paradigm shift, moving from population inferences to the study of individual differences (Dubois & Adolphs, 2016; Seghier & Price, 2018). Previous studies have paved the way for the study of individual variability in functional connectivity patterns of the human brain (Finn et al., 2015; Miranda-Dominguez et al., 2014; Mueller et al., 2013). In this context, resting-state fMRI (rs-fMRI) showed to be particularly powerful in determining underlying differences in the wiring patterns of functional connectome (FC) profiles. Indeed, connectome-based individual predictions achieved identification accuracies as high as 99% when comparing functional connectivity matrices (Finn et al., 2015). Hence, the endeavor to identify and to characterize the individual functional connectivity architecture has been shown to have an imperative place in the study of individual differences.

Recent and mounting evidence suggests that FC profiles are stable at different time scales (Gratton et al., 2018; Jalbrzikowski et al., 2020; Miranda-Dominguez et al., 2018; Sato, White, & Biazoli, 2017). This characteristic posits the FC not only as an individual marker due to the comparably low intrasubject variability but also as a powerful discriminatory measure characterized by the high intersubject variability. Gratton et al. (2018) showed that despite functional networks displaying common organizational features at the group-level, the similarity between functional networks substantially increased at the individual level when evaluating the same participant in different tasks and sessions. This evidence supports the fact that individual stable patterns are crucial for explaining the intersubject variability of functional networks. Therefore, these findings suggest that sources of intersubject variability are stable over time, acting as individual signatures or ‘fingerprints’.

Seghier and Price (2018) refer to the presence of distinct sources of intersubject variability that differ in their timescale. In the lower bound, there are sources of variability due to mood states and context. The medium to long-term sources of intersubject variability include functional patterns built from the intimate interaction of an individual with the environment and genetic factors (Seghier & Price, 2018), respectively. Interestingly, functional networks show distinct levels of intersubject variability. Networks comprising higher-order associative cortical areas seem to remarkably contribute to the FC distinctiveness (Finn et al., 2015; Jalbrzikowski et al., 2020; Kaufmann et al., 2017; Miranda-Dominguez et al., 2018, 2014; Mueller et al., 2013), which, in turn, might be due to a high intersubject (Gratton et al., 2018; Mueller et al., 2013) and low intrasubject variability (Laumann et al., 2015; Poldrack et al., 2015). On the other hand, functional connectivity within networks that comprises primary sensory and motor regions showed high intrasubject and low intersubject variability (Gratton et al., 2018; Laumann et al., 2015; Mueller et al., 2013; Poldrack et al., 2015). The importance of genetic factors to these different levels of intersubject variability, however, is yet to be further investigated.

Recent reports suggest that genetic factors crucially influence the intersubject variability in the functional connectome (Colclough et al., 2017; Demeter et al., 2020; Elliott et al., 2019; Ge, Holmes, Buckner, Smoller, & Sabuncu, 2017; Miranda-Dominguez et al., 2018; Yang et al., 2016). Connectome-based identification analyses were extended to the identification of twin pairs suggesting that part of the intersubject variability is due to genetic factors (Demeter et al., 2020; Miranda-Dominguez et al., 2018). Accordingly, studies indicate that the average heritability of the connectivity strength of the whole-brain connectome is between 15% to 25% within the Human Connectome Project dataset (Adhikari et al., 2018; Colclough et al., 2017; Elliott et al., 2019). On the other hand, the heritability of the connectivity strength within some functional networks seems to be much higher (Ge et al., 2017; Teeuw et al., 2019) than in the whole-brain connectome. However, substantial differences in brain parcellation schemas (Arslan et al., 2018; Eickhoff, Yeo, & Genon, 2018; Salehi et al., 2020) undermines the effort to determine the relationship between heritability and the different levels of intersubject variability. Here, we (1) reproduced and extended the identification analysis introduced by Finn et al. (2015) to determine the functional networks that best uncovered individual uniqueness and intersubject similarity among matched twin pairs, and (2) we investigated how the different levels of intersubject variability of functional networks relate to their heritability by using a multivariate ACE modeling approach with bootstrapping. In our approach, 10 functional connections (edges) were randomly drawn from the pool of connections and were used as variables in a multivariate ACE model. This model decomposes the variance of each variable (i.e., each edge) and the covariance between variables into additive genetic influences (A, or narrow-sense heritability (Mayhew & Meyre, 2017)), shared environment (C) and external sources of variability (E). Here, we only focused on the partitioning of variance to estimate network heritability, doing so by averaging the decomposition of variances into A, C and E components across variables (i.e., across edges) for each model fit. This process was repeated for many iterations which results in the distributions of means for each component (A, C and E). Additionally, this approach allows one to easily generate null distributions for statistical testing by randomly shuffling monozygotic and dizygotic twin statuses at each iteration (Colclough et al., 2017).

## Results

### Functional connectivity-based identification analyses

#### Individual identification

Whole-brain functional connectivity matrices were determined by using two distinct parcellation schemas: “Shen” (Shen, Tokoglu, Papademetris, & Constable, 2013) (268 nodes, 71,824 edges) and “Gordon” (Gordon et al., 2014) (333 nodes, 110,889 edges). For brevity, we only report the results using “Shen” parcels with appropriate reference to equivalent results using “Gordon” parcels in the supplementary material. Connectivity-based identifications were performed comparing pairs of resting-state functional connectivity matrices^4^. Resting-state data were acquired in two different days for every participant included in this study, resulting in two distinct functional connectivity matrices per participant. These pairs of connectivity matrices were separated into a ‘target’ and a ‘database’ set. Individual identification was determined by computing the Pearson’s correlation score of a target connectivity matrix from the ‘target’ set (n=380) with all connectivity matrices from the ‘database’ set (n=380). Following that, the maximum correlation score among all comparisons between the target matrix and each of the FC matrices from the ‘database’ set should correspond to the correlation of the functional connectivity matrices of the same participant in different sessions. This process was repeated for all functional connectivity matrices within the ‘target’ set (Figure 1A). The accuracy of the method was defined by the proportion of correct predicted participants.

**Figure 1.**
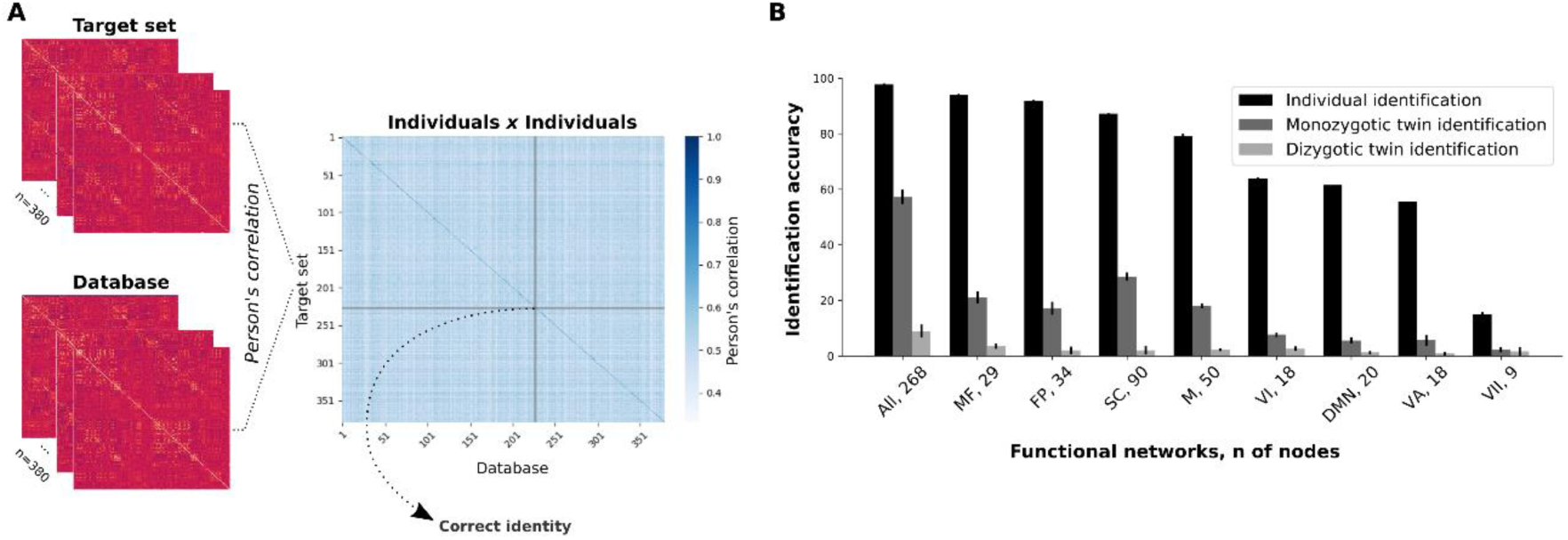
Connectome-based identifications. **A)** Functional connectivity matrices from different sessions were grouped into two datasets, which could be either the ‘target’ set or the ‘database’. Following that, we computed the Pearson’s correlation of each individual connectivity matrix from a ‘target’ set with each connectivity matrix from the ‘database’. Therefore, each row within the individuals vs. individuals matrix contains the correlation scores between a target’s FC and all functional connectivity matrices of the database. **B)** Mean identification accuracies for individual and twin identification analyses for all functional networks (whole-brain included). Mean identification for individual prediction was determined from two combinations of ‘database’ and ‘target’ sets (RESTX × RESTY, where X and Y ∈{1, 2} and X≠Y), while the mean twin identification was determined from four combinations (RESTX × RESTY, where X and Y ∈{1, 2}). Error bars represent the standard deviation. All, whole-brain; MF, medial frontal; FP, frontoparietal; SC, subcortical-cerebellum; M, motor; VI, visual I; DMN, default mode network; VA, visual association; VII, visual II. We also present the number of nodes in each network.

Individual identification analyses were determined with whole-brain functional connectome and individual functional networks (Supplementary Table 1). The resulting accuracy of whole-brain connectome based individual predictions was 97.8% (SD = 0.4%), in agreement with previous studies (Finn et al., 2015; Waller et al., 2017). We also investigated the relevance of individual functional networks for individual predictions by sectioning the whole-brain functional connectome into sub-matrices of single networks. From the 8 functional networks previously defined (Finn et al., 2015), the most successful networks were the medial frontal (93.9 ± 0.5%) and frontoparietal (91.8 ± 0.3%) networks (Figure 1B and Supplementary Table 1). Note that the visual networks and the default mode network were the ones with the worst individual identification accuracy.

#### Twin identification

Previous studies indicate that functional connectivity among higher-order associative brain regions greatly varies across individuals (Gratton et al., 2018; Mueller et al., 2013), even though they are comparably more stable within an individual across sessions (Laumann et al., 2015; Poldrack et al., 2015). Thus, we hypothesized that genetic factors governed sources of high intersubject and low intrasubject variability in the functional connectome. In order to test this hypothesis, we sought to determine whether the FC profiles from pairs of twins were more similar compared to the ones from pairs of unrelated individuals by using connectome-based predictions.

First, we evaluated monozygotic twin identification by computing the correlation coefficients of the functional connectivity matrices of monozygotic individuals (n=246) within the ‘target’ set with all matrices in the database (246×380=93,480 comparisons). Our prediction was based on the selection of the highest correlation score (excluding the correlation scores between functional connectivity matrices of the same individual) for each ‘target’ participant vs. ‘database’ iteration. The mean whole-brain based prediction accuracy was 57.2% (SD = 2.6%). This result indicates that the idiosyncratic FC profiles might be genetically determined and they are sufficiently stable so one could identify monozygotic twins well above chance. Indeed, we have performed a permutation test, by exchanging twin pairs’ identities 1,000 times, such that for each identification iteration, a new twin pair identity was assigned. The maximum identification accuracy found through these 1,000 permutations was 1.6%, indicating that the whole-brain based identification performance is significantly different from the chance level (p-value < 0.001).

Later on, we investigated the ability of specific functional networks in discriminating a twin pair from pairs of unrelated individuals (Figure 1B). At this stage, the most successful functional networks were the subcortical-cerebellum (28.6 ± 1.5%) and medial frontal (21.1 ± 2.2%) networks. Noteworthy, the most successful functional networks on twin identification were amongst the ones that best performed on individual identifications. Nonetheless, a substantial decrease in the successful twin identification rates was observed for functional networks when compared to the whole-brain connectome, and these results were particularly affected by the number of nodes within each network. The least successful functional networks on twin identification were the ones with the least number of nodes, while the networks with a larger number of nodes tended to present higher accuracies. The Pearson’s correlation score between the number of nodes of each network and its ability to correctly identify monozygotic twins was r = 0.95 (p-value = 6.3E-5; Supplementary Table 2), as opposed to a nonsignificant correlation between the number of nodes and individual identification accuracy (r = 0.52, p-value = 0.15). This implies that the ability of *a priori* defined functional networks to capture similarities in the FC profiles of monozygotic twins differentially relies on the amount of information provided (i.e., by the number of nodes).

Finally, we performed all the previous analyses for the identification of dizygotic twins. At this time, we selected only the dizygotic individuals (n=134) within the ‘target’ set, giving 134×380=50,920 comparisons. For the whole-brain based identification, the mean prediction accuracy was 8.9% (SD = 2.3%; p-value <0.001). This abrupt change in twin identification accuracy indicates that the functional connectivity patterns of monozygotic twins are strictly more similar in comparison to dizygotic twins, which indicates the relevance of shared genetic background. At the level of individual functional networks, identification accuracies dropped even further (Figure 1B), and they were also correlated with the number of nodes of the networks (r = 0.92, p-value < 0.001).

### Fingerprinting as a function of the number of edges

The previous results indicated that twin identification accuracy was correlated with the number of nodes of functional networks, and hence with the number of edges. To further investigate the relationship between the number of edges in connectome fingerprinting and twin identification accuracy, we performed identification analyses using randomly selected subsets of edges, with 100 random selections per subset size (Byrge & Kennedy, 2018). Our results show that it is possible to identify an individual with high accuracy using a random subset of edges (Figure 2), with accuracy above 80% using only 500 random edges (a similar finding is reported at Byrge & Kennedy, 2018). However, monozygotic twin identification only reaches near 50% accuracy using 10,000 random edges, while dizygotic twin identification accuracy is equal to 8% on average with the same subset size. Noteworthy, monozygotic twin identification accuracy with 500 random edges was on average equal to approximately 20%, similar to the prediction accuracy using the medial frontal network (29 nodes and 406 unique edges). On the other hand, prediction accuracy reached 32% with 1,000 random edges and 46% with 5,000 random edges. At a similar level, the prediction accuracy of the subcortical-cerebellum network (90 nodes and 4,005 unique edges) was 28.6%.

**Figure 2.**
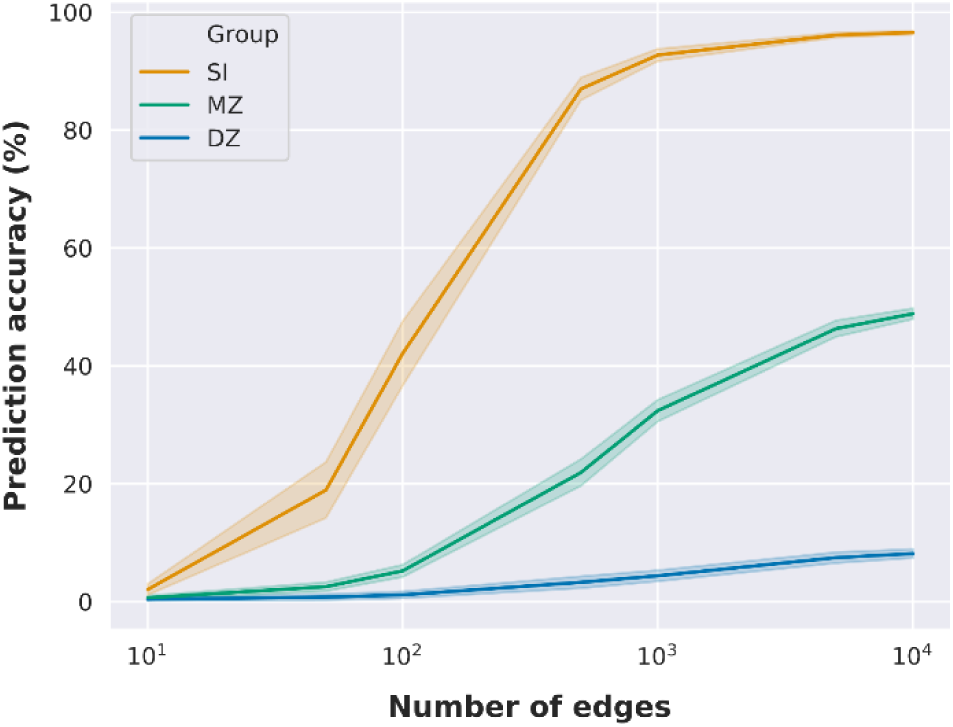
Identification accuracy as a function of the number of edges. Identification accuracy as a function of subsets of randomly selected edges. Mean identification accuracy and standard deviation are illustrated as a function of the number of edges (we only evaluated 7 different subset sizes: 10, 50, 100, 500, 1,000, 5,000, and 10,000 edges). Mean and standard deviation were determined across 100 random edge selections per subset size.

Therefore, our findings suggest that while it is possible to identify twin pairs above chance, differences seen across functional networks in twin pair identification may be mostly driven by differences in the number of nodes/edges. However, the fact that twin identification accuracy with subsets of random edges could outperform functional networks with a similar amount of edges suggests that edges might be differently influenced by genetic factors.

### Intra and intersubject variability in the functional connectome

In order to characterize the intra and intersubject variabilities (i.e. among unrelated individuals, monozygotic and dizygotic twin pairs) for the whole-brain connectome and each functional network, we arranged the correlation coefficients in four groups according to their relationship: 1) same individual - SI (n=380); 2) monozygotic twins - MZ (n=246); 3) dizygotic twins - DZ (n=134) and 4) unrelated individuals - UN (n=143,640). The distributions of correlations across all these pairs for the whole-brain and functional networks are illustrated in Figure 3 (Supplementary Figure 1).

**Figure 3.**
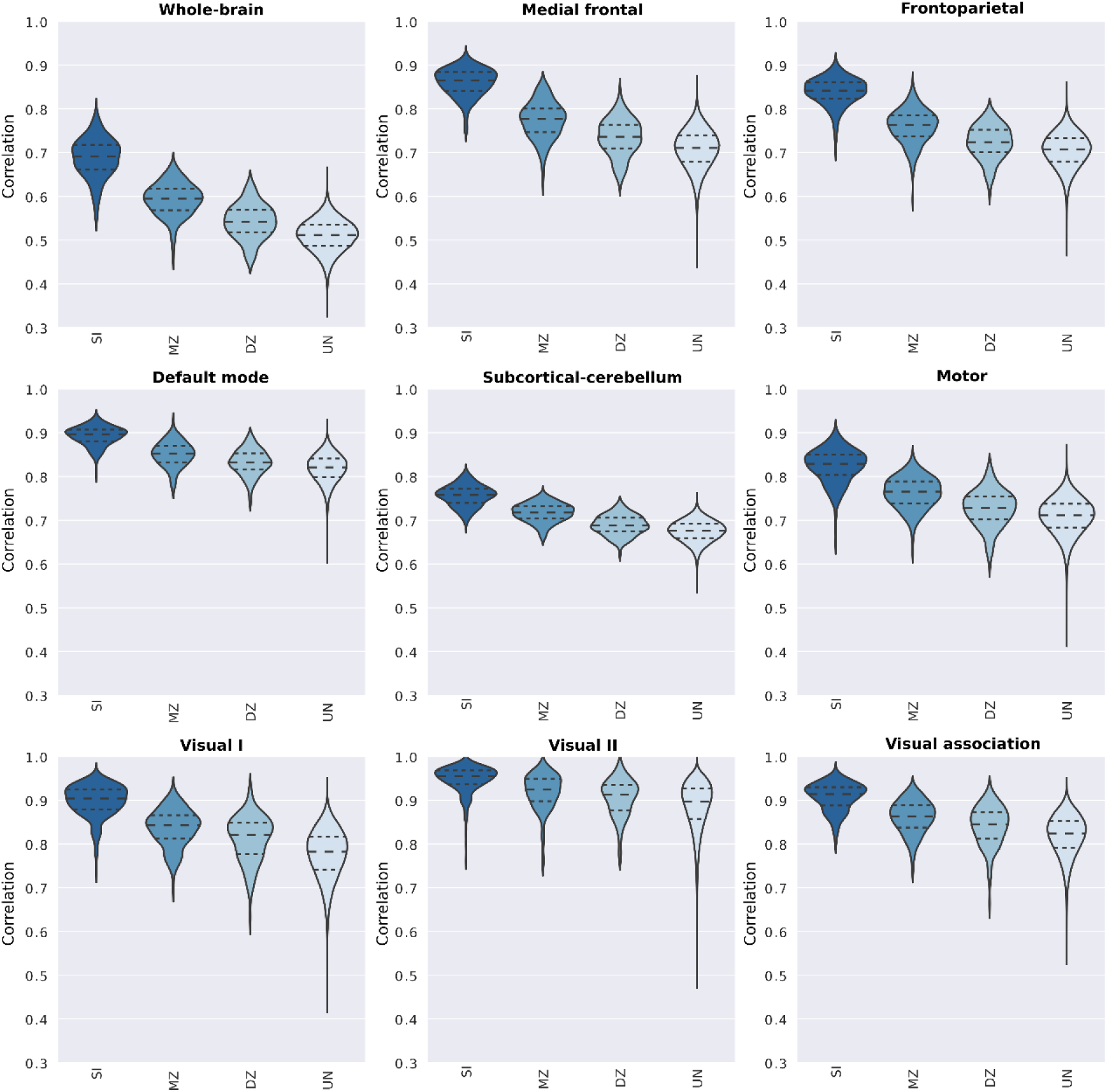
Distribution of correlation coefficients between pairs of functional connectivity matrices for the whole-brain and individual functional networks. Pearson’s correlation scores were determined from pairs of connectivity matrices (REST1 × REST2), and they were grouped based on individuals’ genetic relationship. Hence, violin plots show the distribution of the correlation scores between pairs of matrices of the same individual (SI), monozygotic twin (MZ), dizygotic twin (DZ) and unrelated individuals (UN).

As one could expect, the mean of the distributions of correlation scores from the SI group is notably higher than the ones from the remaining groups. This is observed not only for the whole-brain connectome but also for most of the functional networks, especially for the medial frontal and frontoparietal functional networks. In order to characterize the importance of the distance between these distributions - that is, the effect size - to identification analyses, we determined identification accuracy as a function of effect size, Cliff’s delta (Cliff, 1993) (Figure 4, Supplementary Figure 2 and Supplementary Table 3). In Figure 4, we observe that high prediction accuracy is associated with high effect size, while low prediction accuracy was associated with low effect size. This suggests that high intersubject variability (which is related to low correlation between unrelated individuals’ connectivity matrices) and low intrasubject variability (high correlation between the connectivity matrices of the same individual in different sessions) are crucial for high prediction accuracy. Additionally, the higher similarity between monozygotic twins in comparison to unrelated individuals (medium to high effect sizes) suggests that a portion of this intersubject variability is heritable and differs across functional networks.

**Figure 4.**
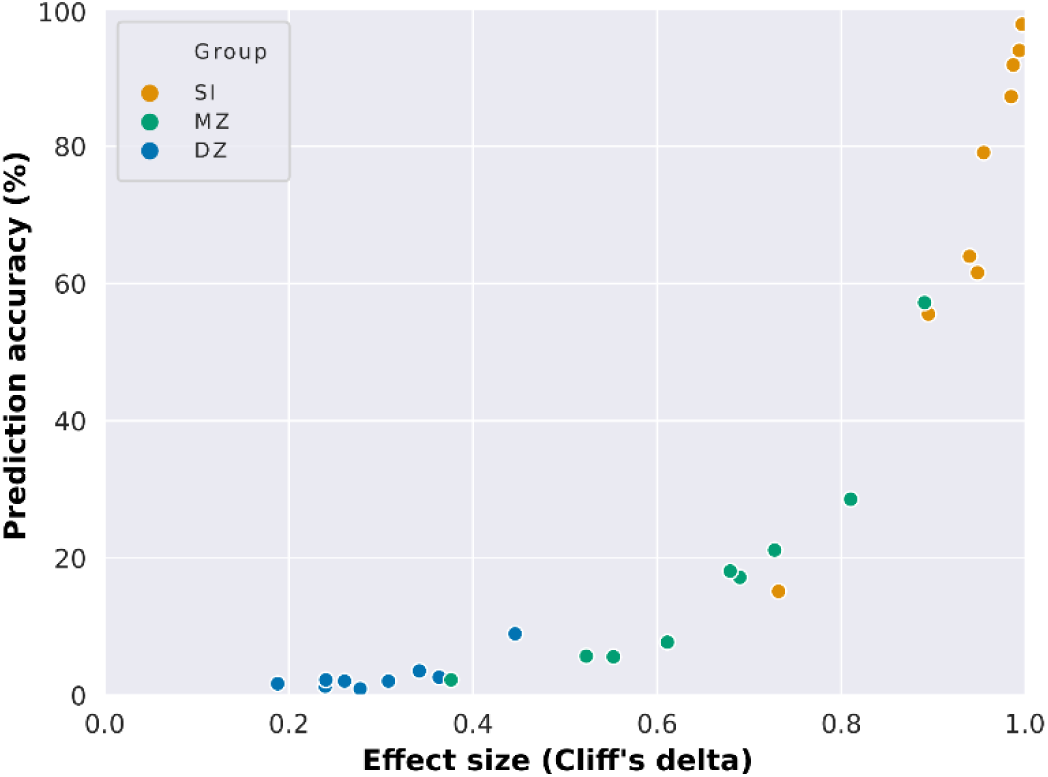
Dependence of connectome-based predictions on effect size. Mean prediction accuracies from all functional networks (whole-brain included) as a function of the effect size of the difference between the group of interest (same individual – SI, monozygotic twins – MZ, or dizygotic twins – DZ) and unrelated individuals.

### Narrow-sense heritability of functional connections

To further investigate these functional networks, we performed heritability analyses using a multivariate ACE modeling approach with bootstrapping. High dimensionality is a common hurdle when multivariate processing is considered for regression or inference methods. Hence, univariate analyses are usually preferred to avoid the necessity of increasing computational resources and time due to high dimensional multivariate analyses trade-off, despite the fact that multivariate analyses tend to be more suitable for complex data that includes several thousand of covariates. In neuroscience, the heritability of functional networks is usually determined as the average heritability of individual functional connections (edges) over their constituent brain regions (nodes) (Colclough et al., 2017; Elliott et al., 2019; Ge et al., 2017). Here, we propose a lower-dimensional multivariate ACE modeling approach with bootstrapping that allows one to generate a distribution of means for each variance component (Figure 5).

**Figure 5.**
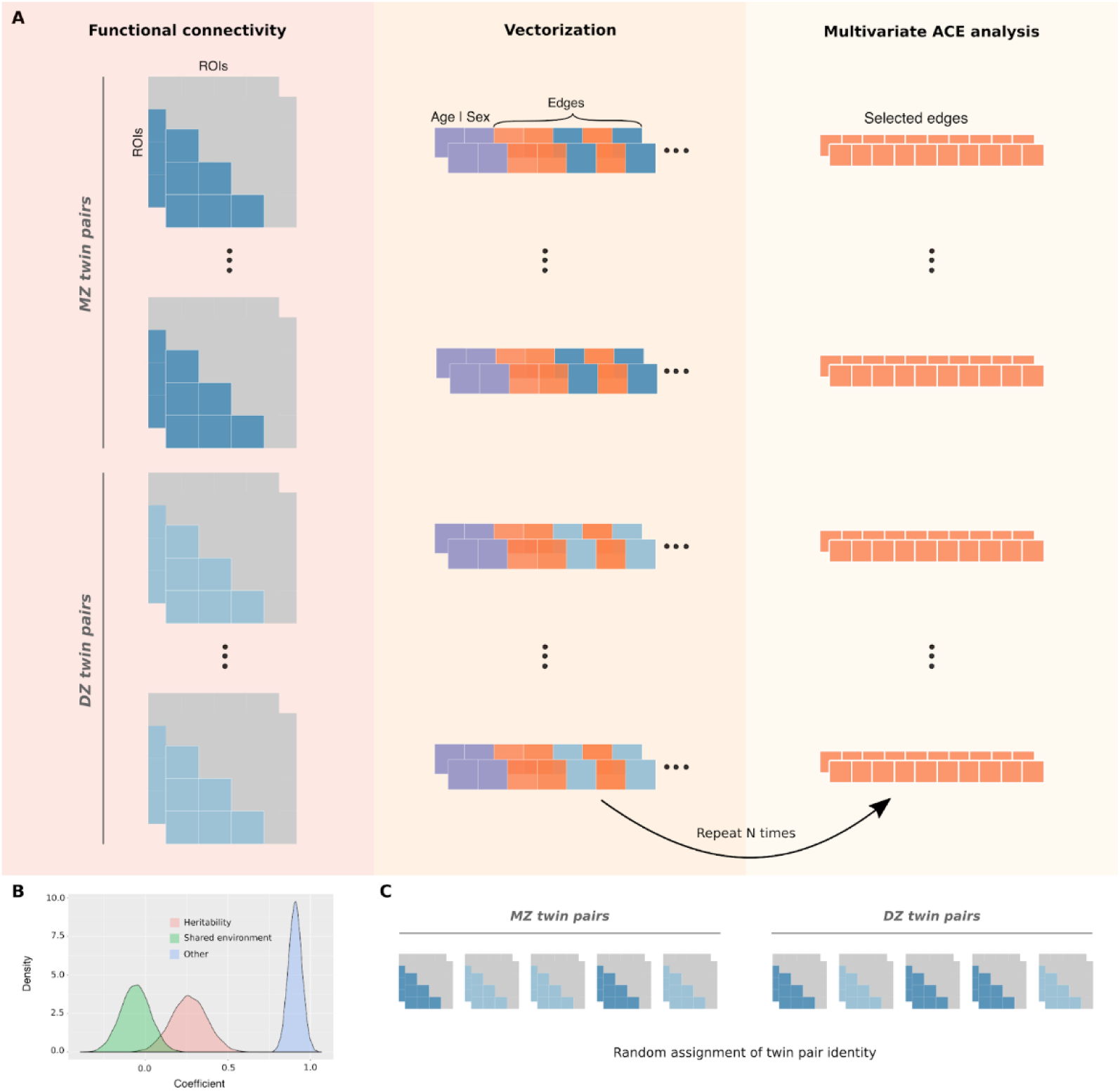
Multivariate ACE model with bootstrapping. **A)** The lower triangles of mean functional connectivity matrices were vectorized, and the effect of age and sex were regressed out from each edge. In an iterative process, 10 edges were randomly selected and used as variables to fit a multivariate ACE model. This procedure was repeated with reposition for 8,000 times for the whole-brain network (or 1,000 times for each functional network). **B)** This approach provides distributions of means for each variance component (A, C and E) by taking the average of the heritability estimates across edges at each iteration. **C)** Null distributions were similarly obtained by randomly shuffling monozygotic and dizygotic twin statuses at each iteration.

This multivariate approach involved the random selection of 10 edges (within the functional network of interest) that were used as variables to fit a multivariate ACE model (Figure 5A). The multivariate ACE model decomposes the variance of each edge into additive genetic influences (A, or narrow-sense heritability (Mayhew & Meyre, 2017)), shared environment (C) and external sources of variability (E). Then, we determined the mean of A, C and E components across edges. This procedure was repeated with reposition for 8,000 times for the whole-brain network and 1,000 times for each functional network, which resulted in the final distributions of means for each component (A, C and E) (Figure 5B). Finally, null distributions were similarly obtained by randomly shuffling monozygotic and dizygotic twin statuses at each iteration (Figure 5C).

The heritability distributions with their respective null distributions for all functional networks are illustrated in Figure 6A (Supplementary Figure 3). As expected, the mean heritability of all null distributions was virtually equal to zero. Apart from that, all heritability estimates distributions were significantly different from their respective null distributions (independent t-test, p<.001). Among all functional networks, the visual II has shown to be the most heritable with mean heritability of 0.37 (37% of the variance of the phenotype is attributed to additive shared genetics; Supplementary Table 4), while the subcortical-cerebellum was the least heritable with mean heritability of 0.20 (Figure 6B and Supplementary Table 4, 5 and 6). Additionally, we compared the mean heritability found for all functional networks using our approach with the mean estimates based on univariate models (Figure 6C). As expected, the mean heritability found using our approach is nearly equal to the classic univariate heritability (Supplementary Table 7), which is based on averaging estimates across all functional connections within each functional network. Finally, heritability estimates were not significantly correlated with number of nodes (r = -0.34, p-value = 0.38) nor monozygotic twin identification accuracy (r = -0.33, p-value = 0.39).

**Figure 6.**
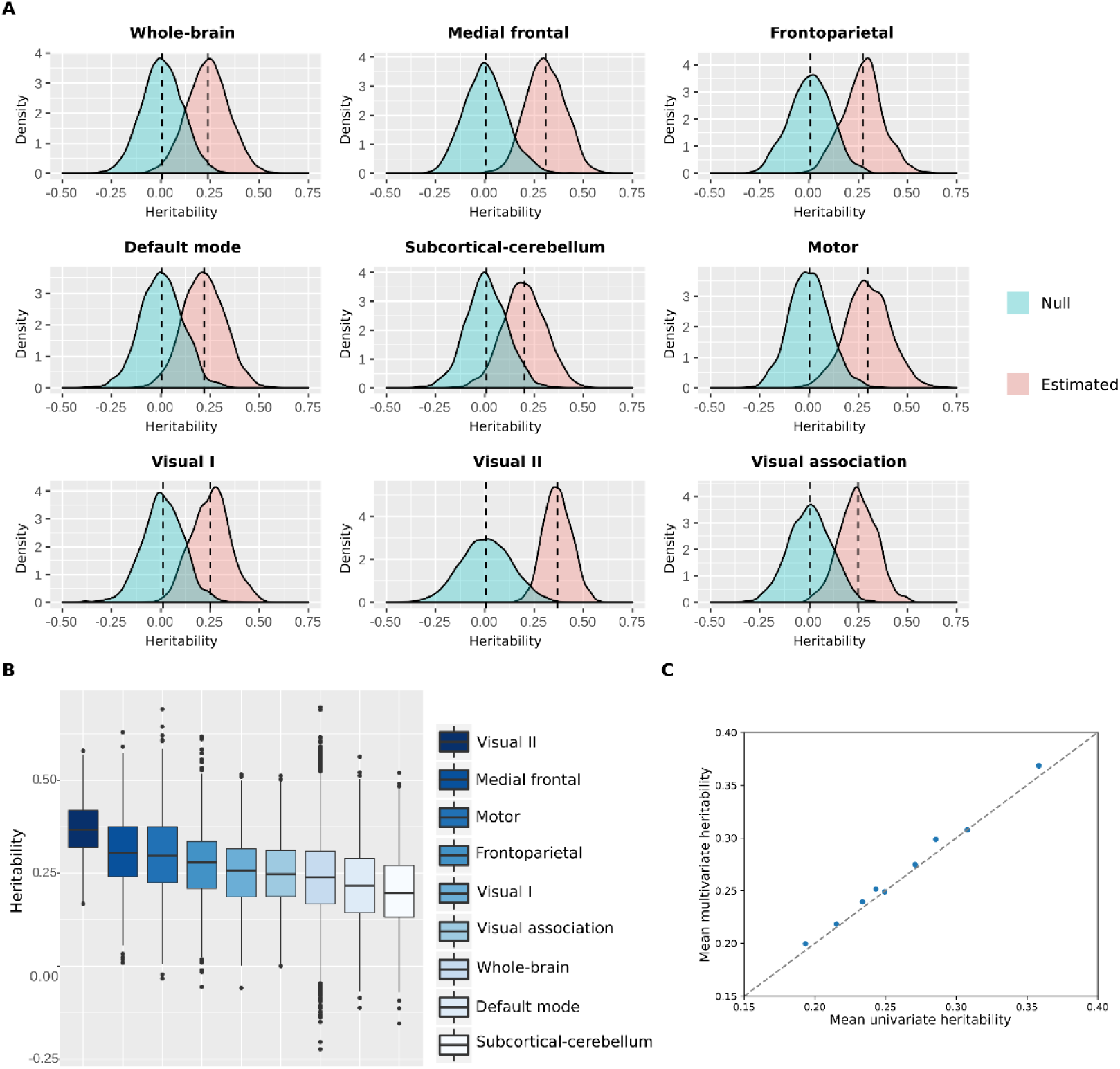
Heritability distributions for each functional network. **A)** Heritability estimates and null distributions for each functional network. **B)** Heritability estimates distributions displayed from the most heritable (visual II) to the least heritable (subcortical-cerebellum). **C)** Comparison of the mean heritability found with multivariate ACE models *versus* univariate ACE models for all functional networks.

## Discussion

Here, we found that the functional connectivity profiles of twin pairs were more similar than of unrelated individuals, although the degree of similarity varied across functional networks. Indeed, we demonstrated that functional networks have distinct discriminatory power in connectome fingerprinting analyses, in both individual and twin identifications, although in the latter differences in identification performances may be mostly driven by differences in the number of nodes/edges. We also found that high intersubject variability (i.e. variability of a trait between individuals) is crucial for connectome fingerprinting. Finally, our multivariate ACE modeling approach suggests that the heritability of functional networks are consistent throughout the brain, although our findings suggest that functional networks are differentially influenced by additive genetic factors. Altogether, we were able to establish the influence of genetic factors to intersubject variability of functional networks by leveraging a multivariate ACE model in addition to the multivariate connectome fingerprinting approach.

### Intra and intersubject variability trade-off in connectome fingerprinting

Evidence suggests that the different levels of inter and intrasubject variability in functional networks contribute to their distinctiveness, such that high intersubject (Gratton et al., 2018; Mueller et al., 2013) and low intrasubject (Laumann et al., 2015; Poldrack et al., 2015) variability in higher-order associative networks are often related to their high discriminability (Finn et al., 2015; Jalbrzikowski et al., 2020; Kaufmann et al., 2017; Miranda-Dominguez et al., 2018, 2014; Mueller et al., 2013) and the opposite pattern to the low discriminability of primary sensory and motor networks (Gratton et al., 2018; Laumann et al., 2015; Mueller et al., 2013; Poldrack et al., 2015). We confirmed that higher-order associative networks were the most discriminatory, while visual networks were the least discriminatory, although they showed similar levels of intrasubject variability. This finding was similarly seen in twin pair identifications, although in the latter the prediction accuracy was positively correlated with the number of nodes defining each functional network. To further investigate the inter and intrasubject variability trade-off in connectome fingerprinting, we determined the prediction accuracy as a function of the difference between the similarity scores of functional networks derived from the same individual - in different resting-state sessions - and unrelated individuals. We found that high identification accuracy requires high intersubject variability, suggesting that although the stability of idiosyncratic functional connectivity patterns is relevant and seen across all functional networks, fingerprinting seems to rely prominently on high intersubject variability.

### Genetic influence on functional networks

To investigate the impact of additive genetic factors in determining stable patterns of intersubject variability, we performed an alternative approach to the univariate ACE model. In our multivariate ACE model, a fixed number of edges were randomly and iteratively selected to fit the model, and the mean heritability estimate was determined by averaging individual edges heritability at each of those iterations. Therefore, 8,000 models were fitted to estimate the heritability of the whole-brain network, as opposed to fitting 35,778 univariate models. In addition to that, 1,000 models were generated for each functional network, totaling 16,000 models (8,000 models for the whole-brain network + 8 * 1,000 models), which is still far less than fitting 35,778 univariate models. We also observed a gain in statistical power with our approach (this is illustrated by the narrower confidence intervals of the multivariate model – Supplementary Table 4 – as opposed to the univariate version – Supplementary Table 7). Additionally, our modeling approach provides a straightforward way for building null distributions by randomly shuffling twin statuses at each iteration as the final step before heritability estimation. Therefore, we believe that the contribution of this method is twofold: it reduces the number of models to be fitted for the estimation of the heritability of functional networks and it also provides a straightforward way for building null distributions.

We found that the functional networks that were the most influenced by additive genetic factors were not the ones that best performed on twin identifications. This is particularly prominent for the visual II and subcortical-cerebellum functional networks. The first has shown to be highly influenced by additive genetic factors, but it had a poor performance on monozygotic twin identification and individual identification. This indicates that the intersubject variability was low, thus being difficult to discriminate between pairs of connectomes from UN/Twin/SI groups. However, a great portion of this low intersubject variability might be due to additive genetic factors. On the other hand, the subcortical-cerebellum network has shown lower heritability but the best performance on twin identification (after whole-brain network). A possible explanation for this finding is that a high intersubject variability allowed a better discrimination between unrelated individuals *versus* twin pairs, even though a smaller portion of its intersubject variability was due to additive genetic factors. Nonetheless, our findings also suggest that twin identification accuracy of functional networks varies with the number of edges, indicating that the inconsistency seen between twin identification accuracy and heritability is perhaps an artefact associated with the confounding effect of number of edges on twin identification.

Finally, heritable patterns of functional connectivity strength of individual edges may emerge from underlying brain anatomy. Anatomical features of the brain have been shown to be highly heritable (Panizzon et al., 2009; Roshchupkin et al., 2016; Strike et al., 2015; Thompson et al., 2001). This suggests that the similarity of brain anatomy in twins might lead to better alignment of their brain structure to a template space as opposed to unrelated individuals. Therefore, when functional units of the brain are determined by a group-based parcellation, variability in functional connectivity strength partly reflects how well a template parcel matches the actual functional unit of a given individual. For example, a given region A in a group-based parcellation could not only overlap with distinct regions across unrelated individuals, but also consistently overlap with a similar area in twins (Anderson et al., 2020). This could lead to the greater similarity of individual edges between twins and higher inter-subject variability across unrelated individuals just because regions being selected are ultimately different. We believe that assessing heritability of functional connectivity patterns using individualized parcellations (Glasser et al., 2016; Kong et al., 2019) might shed some light into this issue.

### Parcellation schema

The individual and twin identification analyses resulted in high prediction accuracy using both parcellation schemas, “Shen” and “Gordon”. Notably, individual identification accuracies using “Shen” parcellation schema is about the same as in previous studies (Finn et al., 2015; Waller et al., 2017), even though we have a more homogenous sample. At the network-level, higher-order associative networks were particularly better at discriminations. This result further supports that associative networks accommodate higher intersubject variability in comparison to sensorimotor networks (Gratton et al., 2018). Despite that, we observed that the default mode network (DMN) defined by both parcellation schemas differed in performance during identification analyses. For “Gordon” parcels, the DMN figured among the most distinctive networks, similarly to other associative networks. However, this pattern was not observed using “Shen” parcels, in which the defined DMN figured among the worst functional networks on individual predictions. This distinction could be due to the different number of nodes attributed to DMN in both schemas. Another finding is that the heritability level of functional networks differed between parcellations, although the mean heritability of the whole-brain functional network was 0.18 using “Gordon” parcels and 0.24 using “Shen” parcels (Supplementary Table 4). This suggests that different brain areas definition greatly impact on heritability estimates, which is a potential topic for further investigation.

Using “Gordon” parcellation, we found that the cingulo parietal and retrosplenial temporal networks were the most influenced by additive genetic factors, whilst the somato-sensory mouth and salience networks were the least ones. On the other hand, Miranda-Dominguez et al. (2018) found that the retrosplenial temporal and somato-sensory mouth were the most heritable, and the visual and salience networks the least heritable. Additionally, their heritability estimates ranged from 0.11 to 0.14, with the heritability of the whole-brain network being equal to 0.20 (Miranda-Dominguez et al., 2018); while our estimates ranged from 0.47 to 0.12. These differences are likely due to differences in heritability estimation approaches; whilst we used the conventional ACE modeling approach, they used three-way repeated-measures ANOVAs. Although the heritability estimates we obtained using “Shen” parcels were more homogeneous, we were still able to capture the different levels of heritability of functional networks, suggesting that our approach is suitable for capturing such differences. Additionally, using a similar methodology, Colclough et al. (2017) found that the heritability of the connectivity strength averaged over parcels was 0.17 for the whole-brain network, and Elliot et al. (2019) found a value of about 0.20. This suggests that, although heritability estimates of functional networks vary depending on the parcellation being used, the whole-brain functional network heritability seems to be reasonably consistent across studies using different methodologies and parcellations.

### Limitations

The effect of head motion on rsfMRI functional connectivity has been assessed over the last decade, and evidence suggests that head motion parameters systematically affect functional connectivity estimates. Van Dijk, Sabuncu, & Buckner (2012) found that increasing mean motion was significantly associated with decreased functional correlation strength among regions in the DMN and the frontoparietal control network, even after regressing out six parameters from the rigid body head motion correction at the preprocessing stage. On the other hand, high levels of head motion were associated with increased local functional connectivity. Finally, their findings suggested that aspects of head motion may behave as trait, which was further investigated by Couvy-Duchesne and colleagues. In Couvy-Duchesne et al. (2014), the influence of additive genetics and environment factors on three head motion parameters have been estimated, and their findings suggest that head motion is partially heritable. These findings effectively suggest that head motion not only systematically affects functional connectivity but it is also partially heritable, indicating that head motion may bias heritability estimates of functional connectivity strength.

The effect of additional preprocessing steps on the confounding effect of head motion in functional connectivity has been systematically investigated (Siegel et al., 2017). Researchers found that extra preprocessing steps to the HCP minimally preprocessed dataset have substantially reduced the correlation of head motion with functional connectivity. Here, we have similarly added extra preprocessing steps to the HCP minimally preprocessed dataset, including CompCor, temporal band-pass filtering, and participants’ movement parameters were used as first-level covariates to regress out their linear components from the BOLD time series. However, it is important to note that complete removal of the spurious effect of motion through regression is difficult (if not impossible). Thus, we believe that the field would benefit from more studies that systematically assess the effect of removing motion parameters at different stages on heritability estimates of functional connectivity.

### Future directions

Our multivariate ACE model suggests that part of the intersubject variability seen in functional networks is due to genetic factors. Transcriptomics and genomics approaches have indicated that many brain disorders are, at least partly, determined by the genetic background (Gandal et al., 2018; Kasten et al., 2018; Prata, Costa-Neves, Cosme, & Vassos, 2019; Sims, Hill, & Williams, 2020). Additionally, disruptions in the human functional and structural connectomes have been associated with neurological conditions, such as amyotrophic lateral sclerosis (ALS) (Chenji et al., 2016), Parkinson’s disease (Gratton et al., 2019; Hall et al., 2019), and epilepsy (Lee et al., 2018). Specifically, neurotoxic accumulation of amyloid plaques in Alzheimer’s disease has been located in areas consistent with cortical hubs, indicating that while cortical hubs are fundamental for information processing, they also bring vulnerability to the human brain (Buckner et al., 2009). Also, many compelling studies have linked psychiatric disorders to fundamental connectome disruptions (van den Heuvel & Sporns, 2019). Despite their unique functional and structural connectivity patterns, these conditions also exhibit some shared patterns that differ from healthy connectomes. The common features of many of these disorders make it difficult to diagnose them and to determine the mechanisms behind their onset, particularly for psychiatric disorders. Thus, detailed scrutiny of the human connectome and genome may lead to a promising new era for precision medicine in psychiatry and neurology. Connectome fingerprinting in addition to heritability analyses may allow for the search of connectome features that bring general and specific vulnerabilities to the human brain, which may be highly heritable, and are central factors among brain disorders (van den Heuvel & Sporns, 2019).

Finally, it is important to acknowledge that although we found differences in heritability estimates across functional networks, such estimates of heritability could be susceptible to different models of heritability. For example, heritability could be better explained with an AE model, in which variance is decomposed into additive genetic factors (A) and external sources of variability (E) only. Additionally, the low reliability of individual edges’ connectivity strength (Noble, Scheinost, & Constable, 2019; Noble et al., 2017) and higher reliability of the connectome as whole suggests that common (shared among edges) and specific (non-shared) sources of genetic variance may differ. The multivariate ACE model used here has been used before to estimate the genetic correlation between two traits, cortical surface area and cortical thickness (Panizzon et al., 2009). However, we believe that a common pathway model would be the most suitable model to study common sources of genetic variance of many edges (Couvy-Duchesne et al., 2014). Therefore, although we found differences in how additive genetic factors may be influencing intersubject variability of functional networks, such estimates are not definite. Critically, different models’ assumptions may potently lead to inconsistent findings of heritability estimates for large-scale functional networks, and future refinements of such estimates (using meta-analysis, for instance) should consider them.

## Materials and Methods

### Database and participant information

In this study, we used the dataset from the “1200 subjects data release” of the Human Connectome Project – HCP (Van Essen et al., 2013). We restricted our analysis to monozygotic (MZ) and dizygotic (DZ) individuals as indicated by genotyping information. So, we initially selected all MZ and DZ individuals from the original sample. From this subsample, we excluded the participants who did not have both resting-state fMRI sessions (ICA-FIX versions) available, and who did not have the twin within the group. Therefore, our final sample size was n=380. Table 1 summarizes the demographic data.

**Table 1.** Demographic information.

|  | Monozygotic<br>(n=246) | Dizygotic<br>(n=134) |
| --- | --- | --- |
| Age, y |  |  |
| Mean ± SD | 29.4 ± 3.3 | 29.1 ± 3.5 |
| Range (min-max) | 22 – 36 | 22 – 35 |
| Sex, n (%) |  |  |
| Female | 144 (58.5) | 78 (58.2) |
| Male | 102 (41.5) | 56 (41.8) |

### Data acquisition

The acquisition protocol has been previously described (Van Essen et al., 2013). In summary, functional and structural data were acquired in a 3T Siemens Skyra scanner using a 32-channel head coil. Resting-state data were collected in two separated sessions (REST1 and REST2) in different days, each session containing two runs of 15 minutes. In this protocol, participants had to keep their eyes open with a relaxed fixation on a projected bright cross-hair in a dark background. Each run within a session is distinguished by the oblique axial acquisition, of which one run used phase encoding in a right-to-left (RL) direction and the other used phase encoding in a left-to-right (LR) direction.

### Data pre-processing

#### Pre-processing pipeline

For this study, we used the spatial and temporal pre-processed rs-fMRI timecourses (Glasser et al., 2013; Smith et al., 2013), which have undergone the steps of artifact removal, motion correction, and registration to standard space. Furthermore, we applied additional pre-processing steps by using the CONN toolbox (v.17.f) (Whitfield-Gabrieli & Nieto-Castanon, 2012), which included: structural segmentation, functional outlier detection (intermediate setting: 5 for z-score scan-to-scan global signal changes and 0.9 mm for scan-to-scan head-motion composite changes), and functional smoothing. Following that, a component-based noise correction method (CompCor) (Behzadi, Restom, Liau, & Liu, 2007) and a temporal band-pass filtering (preserving frequencies between 0.01 and 0.10Hz) were applied. For spatial smoothing, a Gaussian with the full width at half maximum (FWHM) equal to 6mm was used. We also included participant movement parameters as first-level covariates to regress out their linear components from the BOLD time series.

#### Parcellations and functional networks

Timecourses were calculated as the mean signal within the regions of interest (ROIs) defined by different parcellation schemas used: “Gordon” (Gordon et al., 2014) and “Shen” (Shen et al., 2013). Both “Gordon” and “Shen” schemas are data-driven parcellation schemas. The first defines 333 ROIs clustered in 12 functional networks (Supplementary Table 1), in addition to 47 ROIs not assigned to any specific network. The latter defines 268 ROIs clustered in 8 networks (Supplementary Table 1).

#### Functional connectivity matrices

Finally, for the two resting-state sessions, data from both the left-right (LR) and right-left (RL) phase-encoding runs were used to calculate the connectivity matrices. To obtain the connectivity matrices, ROI-to-ROI bivariate correlation connectivity measures were computed for all ROIs defined by both parcellation methods, obtaining two symmetric connectivity matrices for each session for each participant.

### Individual identification

The identification analysis was based on previous work (Finn et al., 2015) with few alterations. Initially, two databases were created containing the functional connectivity matrices for each session (REST1 and REST2). The individual identification was determined by computing the Pearson’s correlation of each individual connectivity matrix from one database with all the other connectivity matrices from the second database (RESTX × RESTY, where X and Y ∈{1, 2} and X≠Y). For a pair of functional connectivity matrices linearly transformed in a column vector (vectorization), T_i_ and D_n_, where T_i_ is the connectivity matrix of a target participant i, and D_n_ is the connectivity matrix of a participant (n=1, …, 380) from the other database, the Pearson’s correlation coefficient r is

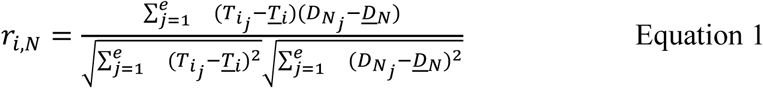

where *e* is the number of edges. In order to predict the identity of the target participant, the maximal Pearson’s correlation coefficient was selected (Figure 1A). Additionally, we also investigated the contribution of single networks to identification accuracy by sub-sectioning the functional connectivity matrices into sub-matrices of single networks. To perform this, we selected only connection within a specified network. Then, we calculated the Pearson’s correlation coefficients, similarly to the previous approach. Results are reported as mean ± SD.

### Twin identification

The twin pair identification algorithm was based on the previous individual identification analysis. At this stage, we removed the correlations corresponding to the same individual in different sessions, that is the diagonal of individuals × individuals matrices, and then performed a new set of identification analyses. In this condition, if the chosen maximum correlation value belonged to the target subject’s twin, the prediction was considered correct. Monozygotic and dizygotic twins were analyzed separately, and all conditions (RESTX × RESTY, where X and Y ∈{1, 2}) were tested. Results are reported as mean ± SD.

### Statistical significance assessment

To assess the statistical significance of twin identification analyses, we performed a permutation testing. To ensure the independence of the dataset, we permuted the twin pairs’ identities, such that for each row of the ‘individuals vs. individuals’ matrix (Figure 1A) a new twin pair identity was assigned. The permutation process was repeated 1,000 times for each functional network.

### Effect size

The distribution of correlation scores between pairs of connectivity matrices (i.e. correlation among the vectorized form of the connectivity matrices) was determined by grouping these scores based on familial relationship: 1) same individual - SI; 2) monozygotic twins - MZ; 3) dizygotic twins - DZ and 4) unrelated individuals - UN. Following that, the effect size of the differences between the distributions of correlation values was measured through the calculation of Cliff’s delta. This a non-parametric effect size measure based on all pairwise differences (Cliff, 1993), which gives how often values from one distribution are larger than the ones from a second distribution (Equation 2).

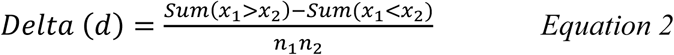

Therefore, the number of times that values from one group are higher than the ones from a second group is calculated for all possible combinations of values between the two groups (n_1_n_2_, where n_1_ and n_2_ are the number of values within the distribution 1 and 2, respectively). The final Cliff’s delta value is the difference between the previous calculations divided by all possible combinations. Thus, a positive and high value of d (dmax_i_mum = 1) mean that values within distribution 1 are mostly higher than the ones within distribution 2, a negative and high absolute value of d (dm_i_n_i_mum = -1) means the other way round, that values within distribution 1 are mostly lower than the ones within distribution 2, and d = 0 means that the distribution 1 and 2 are equal.

### Heritability analyses

Functional connectivity measures from two different days (REST1 and REST2) were averaged, giving a functional connectivity matrix per participant. As mentioned before, whole-brain functional connectivity matrices were determined by using two distinct parcellation schemas: “Shen” (Shen et al., 2013) (268 nodes, 71,824 edges) and “Gordon” (Gordon et al., 2014) (333 nodes, 110,889 edges). The first step involved the vectorization of functional connectivity matrices’ lower triangle (“Shen” - 35,778 edges; “Gordon” – 55,278 edges). The heritability analyses were performed using the umx package (Bates, Maes, & Neale, 2019), after regressing out the effect of age and sex using ‘umx_residualize’.

Heritability of functional networks was estimated using a multivariate ACE model, ‘umxACEv’ from umx package (Bates et al., 2019), with bootstrapping. Specifically, ‘umxACEv’ model allocates observed phenotypic variability of each variable and between variables (variance/covariance matrix) into three latent factors: A (additive genetic factors – h^2^), C (shared environment – c^2^) and E (measurement error or external sources of variability – e^2^) (Neale & Cardon, 1992; Panizzon et al., 2009). This model outputs a variance/covariance load matrix for each component (A, C and E). In each component matrix, the diagonal represents the proportion of variance that that factor explains of each variable’s phenotypic variability, while off-diagonal terms give the proportion of the covariance between variables. Here, we only focused on the partitioning of variance for the estimation of network heritability, doing so by averaging the estimates in the diagonal of each model fit.

In each iteration of model fitting, a subset of 10 edges was randomly selected and used to fit the previously described ACE model. This procedure was repeated with reposition for 8,000 times (or 12,000 times when “Gordon” parcels was used) for whole-brain, and 1,000 times for each functional network. The number of iterations was determined such that every edge would be selected at least twice (i.e. 8,000 iterations * 10 edges = 80,000). This approach provides distributions of means of each component (A, C and E) for each functional network. Finally, null distributions were similarly obtained by randomly shuffling monozygotic and dizygotic twin statuses at each iteration (Colclough et al., 2017). Independent t-student tests were performed separately to evaluate whether each functional network’s heritability distribution significantly differed from their respective null distribution.

### Code availability

All source codes will be available at https://github.com/felenitaribeiro/fingerprinting_twinStudy and https://github.com/frcsantos/heritability upon publication of this manuscript.

### Citation diversity statement

Recent work in neuroscience and other fields identified a bias in citation practices such that papers from women and other minorities are under-cited relative to the number of such papers in the field (Caplar, Tacchella, & Birrer, 2017; Dion, Sumner, & Mitchell, 2018; Dworkin et al., 2020; Maliniak, Powers, & Walter, 2013; Mitchell, Lange, & Brus, 2013). Here we sought to proactively consider choosing references that reflect the diversity of the field in thought, form of contribution, gender, and other factors. Gender of the first and last author of each reference was predicted by using databases that store the probability of a name being carried by a man or a woman (Dworkin et al., 2020). By this measure (and excluding self-citations to the first and last authors of our current paper), our references contain 10.31% woman(first)/woman(last), 18.36% man/woman, 21.55% woman/man, and 49.78% man/man. We look forward to future work that could help us to better understand how to support equitable practices in science.

## Acknowledgments

This work was supported by the Universidade Federal do ABC (UFABC) and Coordination of Improvement of Higher Education Personnel (CAPES). Data were provided by the Human Connectome Project, WU-Minn Consortium (Principal Investigators: David Van Essen and Kamil Ugurbil; 1U54MH091657) funded by the 16 NIH Institutes and Centers that support the NIH Blueprint for Neuroscience Research; and by the McDonnell Center for Systems Neuroscience at Washington University.

## Author contributions

F.L.R., F.R.C.S and C.E.B. conceptualized the study. F.L.R. and F.R.C.S. designed and performed the analyses with support from W.H.L.P. F.L.R. wrote the original draft. All authors revised and edited the manuscript. J.R.S., W.H.L.P., and C.E.B provided support and guidance with data interpretation.

