## Supplementary Material for "Inferring the heritability of large-scale functional networks with a multivariate ACE modeling approach"

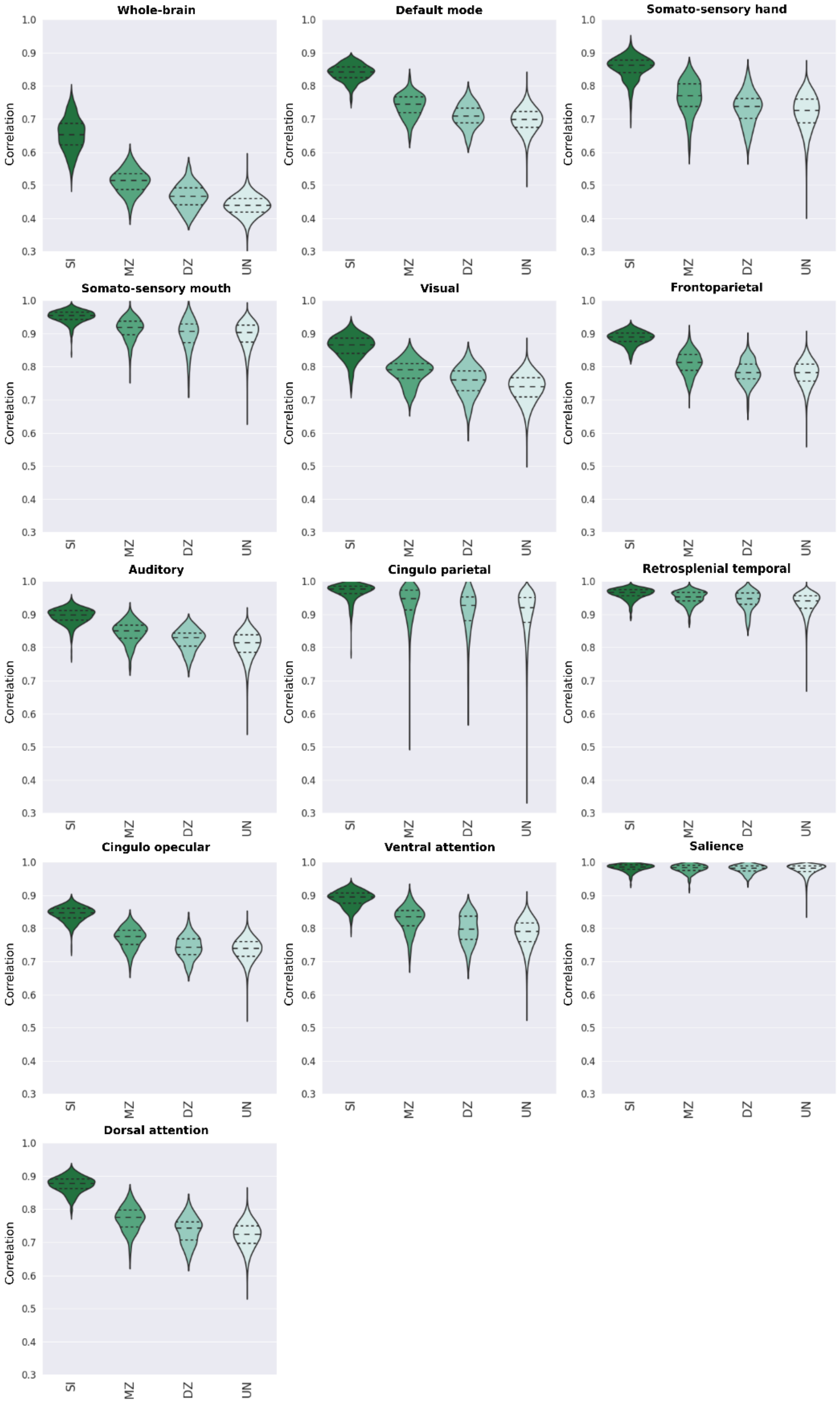


**Supplementary Figure 1 - Distribution of correlation coefficients between pairs of functional connectivity matrices for the whole-brain and individual functional networks defined for "Gordon" parcellation schema.** Pearson's correlation scores were determined from pairs of connectivity matrices (REST1 × REST2), and they were grouped based on individuals' genetic relationship. Hence, violin plots show the distribution of the correlation scores between pairs of matrices of the same individual (SI), monozygotic twin (MZ), dizygotic twin (DZ) and unrelated individuals (UN).

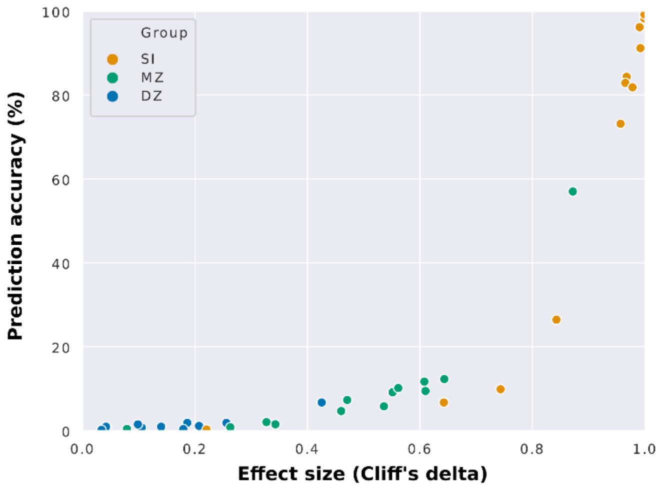


**Supplementary Figure 2 - Dependence of connectome-based predictions on effect size using "Gordon" parcellation.** Here, the mean prediction accuracies from all functional networks (whole-brain included) were plotted as a function of the effect size of the difference between the group of interest (same individual – SI, monozygotic twins – MZ, or dizygotic twins – DZ) and unrelated individuals.

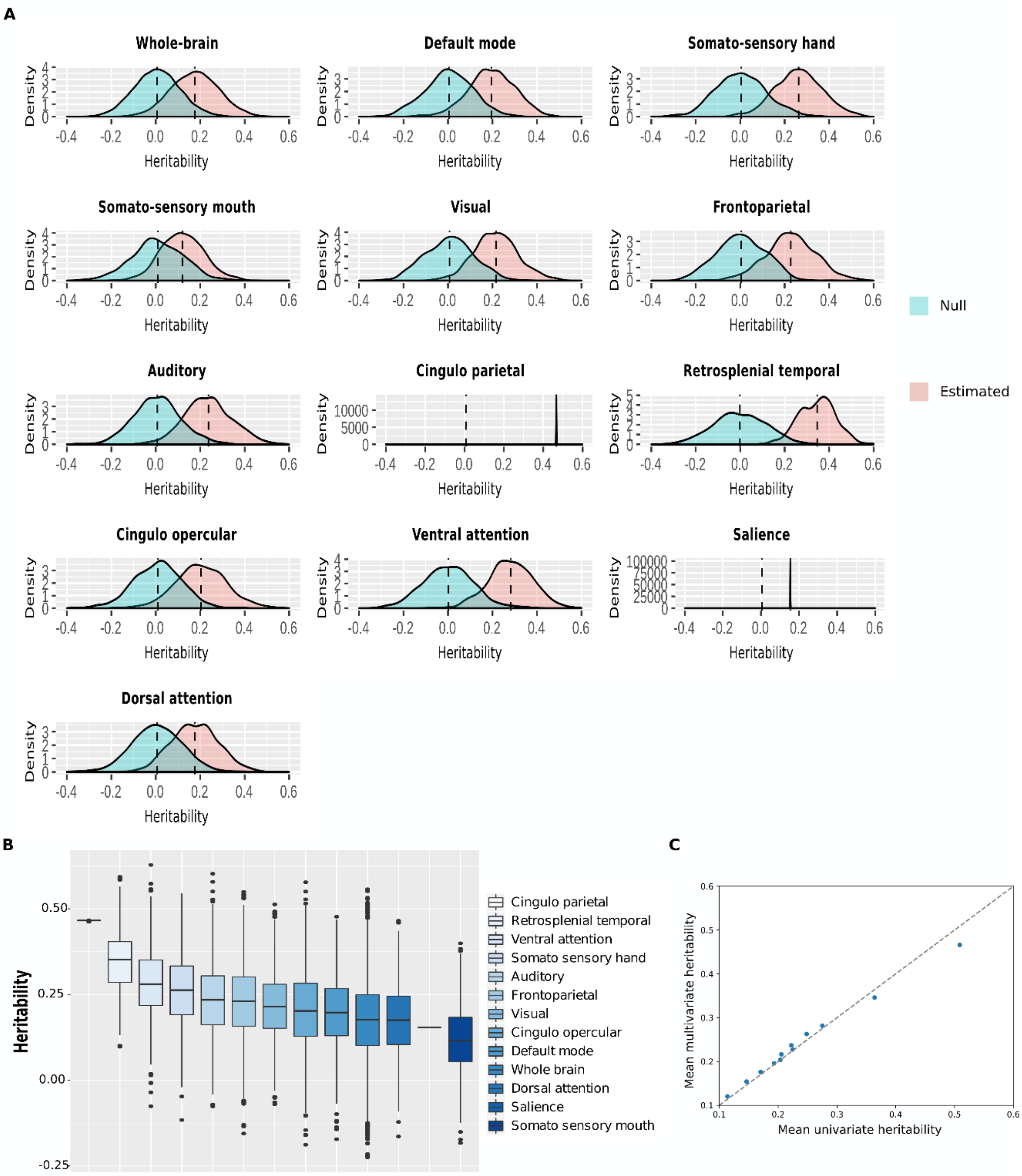


**Supplementary Figure 3 - Heritability distributions for each functional network using 'Gordon' parcellation. A)** Heritability estimates and null distributions for each functional network. **B)** Heritability estimates distributions are shown from the most heritable (cingulo parietal) to the least heritable (somato sensory mouth). **C)** Comparison of the mean heritability found with multivariate ACE models versus univariate ACE models for all functional networks.

**Supplementary Table 1 - Network-based individual and twin identification accuracies using "Gordon" and "Shen" parcellation schemas.** Note: Functional networks were *a priori* defined. For "Gordon" parcellation schema, 47 ROIs were not assigned to any specific functional network. Thus, these ROIs were only considered for whole-brain functional connectivity based identifications. Mean identification accuracies for individual and twin identification analyses for all functional networks (whole-brain included). Mean identification for individual prediction was determined from two combinations of 'database' and 'target' sets (RESTX × RESTY, where X and Y ∈ {1, 2} and X≠Y), while the mean twin identification was determined from four combinations (RESTX × RESTY, where X and Y ∈ {1, 2}).

| | Prediction accuracy ± SD (%) | | | | | |
|---|---|---|---|---|---|---|
| Network, number of ROIs | Individual | | Monozygotic twin | | Dizygotic twin | |
| | Mean | Std | Mean | Std | Mean | Std |
| "Gordon" | | | | | | |
| Whole-brain, 333 | 99.87 | 0.13 | 57.01 | 0.83 | 6.72 | 1.40 |
| Default mode, 41 | 98.29 | 0.13 | 11.69 | 2.50 | 1.87 | 1.24 |
| Smhand, 38 | 84.34 | 0.39 | 7.32 | 1.11 | 0.75 | 0.00 |
| Smmouth, 8 | 26.45 | 0.39 | 0.81 | 0.29 | 0.37 | 0.37 |
| Visual, 39 | 82.89 | 0.53 | 9.45 | 0.88 | 1.87 | 0.65 |
| Frontoparietal, 24 | 91.18 | 1.45 | 4.67 | 0.61 | 0.75 | 0.53 |
| Auditory, 24 | 73.16 | 0.53 | 5.79 | 0.34 | 0.00 | 0.00 |
| Cingulo Parietal, 5 | 9.87 | 0.66 | 2.03 | 1.22 | 0.93 | 0.62 |
| Retrosplenial Temporal, 8 | 6.71 | 0.66 | 1.52 | 0.67 | 0.37 | 0.65 |
| Cingulo Opercular, 40 | 96.18 | 0.39 | 9.15 | 1.09 | 1.49 | 1.06 |
| Ventral Attention, 23 | 81.84 | 0.53 | 10.16 | 0.81 | 0.93 | 0.81 |
| Salience, 4 | 0.26 | 0.00 | 0.41 | 0.70 | 0.19 | 0.32 |
| Dorsal Attention, 32 | 99.21 | 0.00 | 12.30 | 2.31 | 1.12 | 0.65 |
| *"Shen"* | | | | | | |
| Whole-brain, 268 | 97.76 | 0.39 | 57.22 | 2.61 | 8.96 | 2.36 |
| Medial frontal, 29 | 93.95 | 0.53 | 21.14 | 2.23 | 3.54 | 0.97 |
| Frontoparietal,34 | 91.84 | 0.26 | 17.17 | 2.36 | 2.05 | 1.33 |
| Default mode, 20 | 61.58 | 0.00 | 5.59 | 1.20 | 1.31 | 0.62 |
| Subcortical-cerebellum, 90 | 87.24 | 0.13 | 28.56 | 1.50 | 2.05 | 1.53 |
| Motor, 50 | 79.08 | 0.92 | 18.09 | 0.93 | 2.24 | 0.53 |
| Visual I, 18 | 63.95 | 0.26 | 7.72 | 0.86 | 2.61 | 0.83 |
| Visual II, 9 | 15.13 | 0.66 | 2.24 | 0.93 | 1.68 | 1.62 |
| Visual association, 18 | 55.53 | 0.00 | 5.69 | 1.97 | 0.93 | 0.62 |

**Supplementary Table 2 - Correlation score between the number of nodes and the average identification scores.**

| | Pearson correlation | Significance (p-value) |
|---|---|---|
| "Gordon" | | |
| SI | 0.40 | 0.17 |
| MZ | 0.99 | 5.87E-10 |
| DZ | 0.97 | 7.66E-8 |
| "Shen" | | |
| SI | 0.52 | 0.15 |
| MZ | 0.95 | 6.27E-5 |
| DZ | 0.92 | 3.91E-4 |

**Supplementary Table 3 - Effect size of the difference among groups of correlation scores using Cliff's delta.**

| | Cliff's delta | | |
|---|---|---|---|
| | SI vs UN | MZ vs UN | DZ vs UN |
| "Gordon" | | | |
| Whole-brain | 1.00 | 0.87 | 0.43 |
| Default mode | 1.00 | 0.61 | 0.19 |
| Smhand | 0.97 | 0.47 | 0.11 |
| Smmouth | 0.84 | 0.26 | 0.04 |
| Visual | 0.97 | 0.61 | 0.26 |
| Frontoparietal | 0.99 | 0.46 | 0.04 |
| Auditory | 0.96 | 0.54 | 0.21 |
| Cingulo Parietal | 0.74 | 0.33 | 0.04 |
| Retrosplenial Temporal | 0.64 | 0.34 | 0.18 |
| Cingulo Opercular | 0.99 | 0.55 | 0.10 |
| Ventral Attention | 0.98 | 0.56 | 0.14 |
| Salience | 0.22 | 0.08 | 0.03 |
| Dorsal Attention | 1.00 | 0.64 | 0.21 |
| "Shen" | | | |
| Whole-brain | 1.00 | 0.89 | 0.45 |
| Medial frontal | 0.99 | 0.73 | 0.34 |
| Frontoparietal | 0.99 | 0.69 | 0.26 |
| Default mode | 0.95 | 0.55 | 0.24 |
| Subcortical-cerebellum | 0.98 | 0.81 | 0.31 |
| Motor | 0.95 | 0.68 | 0.24 |
| Visual I | 0.94 | 0.61 | 0.36 |
| Visual II | 0.73 | 0.38 | 0.19 |
| Visual association | 0.89 | 0.52 | 0.28 |

**Supplementary Table 4 – Mean heritability (multivariate ACE model) and confidence interval of each functional network.**

| Network | Mean | Lower bound (2.5 percentile) | Upper bound (97.5 percentile) |
|---|---|---|---|
| "Gordon" | | | |
| Whole-brain | 0.18 | -0.04 | 0.39 |
| Default mode | 0.20 | -0.02 | 0.39 |
| Smhand | 0.26 | 0.06 | 0.47 |
| Smmouth | 0.12 | -0.06 | 0.31 |
| Visual | 0.22 | 0.03 | 0.42 |
| Frontoparietal | 0.23 | 0.00 | 0.45 |
| Auditory | 0.24 | 0.03 | 0.45 |
| Cingulo Parietal | 0.47 | 0.47 | 0.47 |
| Retrosplenial Temporal | 0.35 | 0.19 | 0.50 |
| Cingulo Opercular | 0.20 | -0.02 | 0.44 |
| Ventral Attention | 0.28 | 0.08 | 0.47 |
| Salience | 0.15 | 0.15 | 0.15 |
| Dorsal Attention | 0.18 | -0.01 | 0.38 |
| "Shen" | | | |
| Whole-brain | 0.24 | 0.03 | 0.45 |
| Medial frontal | 0.31 | 0.12 | 0.49 |
| Frontoparietal | 0.27 | 0.08 | 0.47 |
| Default mode | 0.22 | 0.02 | 0.43 |
| Subcortical-cerebellum | 0.20 | -0.01 | 0.40 |
| Motor | 0.30 | 0.08 | 0.52 |
| Visual I | 0.25 | 0.07 | 0.45 |
| Visual II | 0.37 | 0.24 | 0.51 |
| Visual association | 0.25 | 0.07 | 0.42 |

**Supplementary Table 5 - Independent t-test p-values of comparisons among functional networks as defined "Shen" parcellation.** Note: Alpha-level corrected for multiple comparisons is equal to 0.0015625.

| | Whole-brain | Medial frontal | Frontoparietal | Default mode | Subcortical-cerebellum | Motor | Visual I | Visual II | Visual association |
|---|---|---|---|---|---|---|---|---|---|
| Whole-brain | | 7.29E-81 | 4.46E-23 | 4.51E-09 | 1.57E-28 | 5.54E-60 | 0.000613 | 6.3E-282 | 0.006782 |
| Medial frontal | | | 1.86E-13 | 1.61E-80 | 3E-111 | 0.055264 | 1.56E-37 | 3.92E-55 | 1.84E-42 |
| Frontoparietal | | | | 2.75E-33 | 1.35E-55 | 6.16E-07 | 1.58E-07 | 1.1E-111 | 2.77E-09 |
| Default mode | | | | | 6.11E-05 | 2.75E-58 | 1.72E-13 | 1.1E-236 | 3.92E-12 |
| Subcortical-cerebellum | | | | | | 4.13E-84 | 7.06E-30 | 3E-278 | 3.08E-28 |
| Motor | | | | | | | 1.28E-23 | 6.83E-59 | 6.06E-27 |
| Visual I | | | | | | | | 1.6E-174 | 0.535596 |
| Visual II | | | | | | | | | 1.2E-189 |
| Visual association | | | | | | | | | |

**Supplementary Table 6 - Independent t-test p-values of comparisons among functional networks as defined "Gordon" parcellation.** Note: Alpha-level corrected for multiple comparisons is equal to 0.05/78=0.000641026.

| | Whole-brain | Default mode | Somato-sensory hand | Somato-sensory mouth | Visual | Frontoparietal | Auditory | Cingulo Parietal | Retrosplenial Temporal | Cingulo Opercular | Ventral Attention | Salience | Dorsal Attention |
|---|---|---|---|---|---|---|---|---|---|---|---|---|---|
| Whole-brain | | 1.14E-08 | 2.2E-130 | 1.37E-55 | 1.16E-30 | 1.16E-47 | 4.75E-66 | 0 | 0 | 4.21E-15 | 1.9E-192 | 0.527916 | 0.919909 |
| Default mode | | | 2.02E-43 | 3.03E-60 | 6.94E-06 | 1.01E-10 | 9.91E-18 | 0 | 9.3E-217 | 0.112817 | 1.02E-72 | 0.206359 | 2.22E-05 |
| Somato-sensory hand | | | | 1.8E-177 | 2.06E-23 | 9.84E-13 | 7.23E-08 | 0 | 2.66E-78 | 1.03E-31 | 3.81E-05 | 0.001319 | 9.58E-70 |
| Somato-sensory mouth | | | | | 8.94E-98 | 5E-104 | 2.2E-126 | 0 | 0 | 2.35E-65 | 3.4E-229 | 0.260407 | 1.42E-34 |
| Visual | | | | | | 0.0189 | 8.93E-06 | 0 | 1.2E-181 | 0.008015 | 7.98E-47 | 0.046014 | 1.42E-18 |
| Frontoparietal | | | | | | | 0.056683 | 0 | 3.4E-136 | 3.02E-06 | 3.16E-29 | 0.039849 | 1.58E-25 |
| Auditory | | | | | | | | 0 | 4.8E-125 | 2.95E-11 | 9.33E-22 | 0.014828 | 2.16E-36 |
| Cingulo Parietal | | | | | | | | | 0 | 0 | 0 | 0 | 0 |
| Retrosplenial Temporal | | | | | | | | | | 3.9E-181 | 1.11E-52 | 2.84E-13 | 3.5E-262 |
| Cingulo Opercular | | | | | | | | | | | 7.41E-56 | 0.171995 | 1.98E-08 |
| Ventral Attention | | | | | | | | | | | | 5.61E-05 | 3E-105 |
| Salience | | | | | | | | | | | | | 0.506939 |
| Dorsal Attention | | | | | | | | | | | | | |

**Supplementary Table 7 – Mean heritability (univariate ACE model) and confidence interval of each functional network**

| Network | Mean | Lower bound (2.5 percentile) | Upper bound (97.5 percentile) |
|---|---|---|---|
| "Gordon" | | | |
| Whole-brain | 0.17 | -0.46 | 0.81 |
| Default mode | 0.19 | -0.46 | 0.83 |
| Smhand | 0.25 | -0.32 | 0.86 |
| Smmouth | 0.11 | -0.65 | 0.87 |
| Visual | 0.21 | -0.38 | 0.81 |
| Frontoparietal | 0.22 | -0.40 | 0.94 |
| Auditory | 0.22 | -0.39 | 0.87 |
| Cingulo Parietal | 0.51 | 0.09 | 0.92 |
| Retrosplenial Temporal | 0.36 | -0.27 | 0.90 |
| Cingulo Opercular | 0.20 | -0.47 | 0.88 |
| Ventral Attention | 0.28 | -0.35 | 0.86 |
| Salience | 0.15 | -0.15 | 0.61 |
| Dorsal Attention | 0.17 | -0.48 | 0.81 |
| "Shen" | | | |
| Whole-brain | 0.23 | -0.40 | 0.86 |
| Medial frontal | 0.31 | -0.27 | 0.82 |
| Frontoparietal | 0.27 | -0.33 | 0.88 |
| Default mode | 0.22 | -0.40 | 0.85 |
| Subcortical-cerebellum | 0.19 | -0.43 | 0.85 |
| Motor | 0.29 | -0.33 | 0.92 |
| Visual I | 0.24 | -0.32 | 0.76 |
| Visual II | 0.36 | -0.16 | 0.79 |
| Visual association | 0.25 | -0.28 | 0.86 |
